# Time series metagenomic sampling of the Thermopyles, Greece, geothermal springs reveals stable microbial communities dominated by novel sulfur-oxidizing chemoautotrophs

**DOI:** 10.1101/2020.07.20.211532

**Authors:** A. Meziti, E. Nikouli, J.K. Hatt, K. Konstantinidis, K. Ar. Kormas

## Abstract

Geothermal springs are barely affected by environmental conditions aboveground as they are continuously supplied with subsurface water with little variability in chemistry. Therefore, changes in their microbial community composition and function, especially over a long period, are expected to be limited but this assumption has not yet been rigorously tested. Toward closing this knowledge gap, we applied whole metagenome sequencing to 17 water samples collected between 2010 and 2016 (two to four samples per year) from the Thermopyles sulfur geothermal springs in central Greece. As revealed by 16S rRNA gene fragments recovered in the metagenomes, *Epsilonproteobacteria*-related operational taxonomic units (OTUs) dominated most samples, while grouping of samples based on OTU abundances exhibited no apparent seasonal pattern. Similarities between samples regarding functional gene content were high, especially in comparison to other surface water systems in Greece, with all samples sharing >70% similarity in functional pathways. These community-wide patterns were further confirmed by analysis of metagenome-assembled genomes (MAGs), which showed - in addition- that novel species and genera of the chemoautotrophic *Campylobacterales* order dominated the springs. These MAGs carried different pathways for thiosulfate and/or sulfide oxidation coupled to carbon fixation pathways. Overall, our study showed that even in the long term, functions of microbial communities in a moderately hot terrestrial spring remain stable, driving presumably the corresponding stability in community structure.

## INTRODUCTION

Microbial communities of geothermal springs are of special interest for understanding early life as they are considered analogues of the first habitable environments on Earth (Damer & Deamer, 2019; Konhauser, Phoenix, Bottrell, Adams, & Head, 2001). These habitats are dominated by several thermophilic and hyperthermophilic species (López-López, Cerdán, & González-Siso, 2013) that mediate nutrient cycling (e.g., C, N, S) (Falkowski, Fenchel, & Delong, 2008). The best-studied geothermal habitat is the Yellowstone National Park (USA) which contains >14,000 of sites with geothermal activity, covering a wide range of pH (2–10), temperature (40–92°C), and geochemical properties (Rye & Truesdell, 2007). Most of these sites exhibit temporal stability in these geochemical properties (Inskeep et al., 2013).

Several molecular studies of the microbial diversity present in Yellowstone Park have revealed the dominance of uncultivated thermophilic archaea, *Aquificales* and *Cyanobacteria* at different sites (Barns, Fundyga, Jeffries, & Pace, 1994;Barns, Delwiche, Palmer, & Pace, 1996; Boomer, Lodge, Dutton, & Pierson, 2002; Hugenholtz, Pitulle, Hershberger, & Pace, 1998; Meyer-Dombard, Shock, & Amend, 2005; Reysenbach et al., 2006; Toplin, Norris, Lehr, McDermott, & Castenholz, 2008). Subsequent studies, including a large metagenomic survey (Inskeep et al., 2013), tried to elucidate the geochemical features that drive microbial diversity patterns and how specific phylotypes are functionally differentiated from each other based on electron donors such as hydrogen or sulfide and electron acceptors. Other habitats that have been studied, including thermophilic springs in Japan (Yamamoto et al., 1998), Iceland (Flores, Liu, Ferrera, Beveridge, & Reysenbach, 2008;Takacs-Vesbach, Mitchell, Jackson-Weaver, & Reysenbach, 2008; Tobler & Benning, 2011), New Zealand (Childs, Mountain, O’Toole, & Stott, 2008) and Kamchatka (Russia; Burgess, Unrine, Mills, Romanek, & Wiegel, 2012; (Wilkins, Ettinger, Jospin, & Eisen, 2019) were found to harbor similar microbial communities. Most of these habitats exhibit high temperatures (>65 °C), and thus mainly consist of hyperthermophilic communities.

Mesophilic sulfur-rich terrestrial springs, in particular, are comparatively much less studied and understood. Chemoautotrophic microbial communities use reduced sulfur compounds contained in underlying geothermal water to gain energy and fuel these systems. These communities are important in sulfur cycling and carbon fixation, thus mediating the transfer of energy from the geothermal source to other trophic levels (Hügler, Gärtner, & Imhoff, 2010). Most studies of mesophilic terrestrial springs have analyzed microbial community diversity and/or functions in order to detect key community members and the functions performed in association with environmental and/or geothermal parameters (Chan, Chan, Tay, Chua, & Goh, 2015; Engel, Porter, Stern, Quinlan, & Bennett, 2004;, Headd & Engel, 2013; Reigstad, Jorgensen, Lauritzen, Schleper, & Urich, 2011). However, although steady geochemical conditions imply stability in microbial diversity and function, no study has been performed on a temporal or seasonal scale. Temporal analysis is particularly important for mesophilic springs since these systems typically receive soil or water inputs from other sources (i.e., seawater or rainfall), in addition to groundwater. It currently remains unknown whether these inputs could influence major metabolic pathways and override the major input from geothermal fluids.

Greece harbors many terrestrial springs due to the geology of the country. The formation of terrestrial springs is related to recent volcanic activity and active tectonics for which magmatic and volcanic processes along with the high mountain chains and active fault systems favor the rise of deep waters discharged at the surface as geothermal springs. Most springs are characterized by the mixing of deep thermal reservoir water with meteoric water, while coastal springs are also characterized by the mixing of geothermal water with seawater and/or freshwater, and thus losing their heat.

The Thermopyles springs located in the eastern part of mainland Greece (38°47’36.35“N/ 22°31’43.04”E) consist of such typical coastal springs and comprise one of the larger active geothermal systems in Greece. They are part of the Spercheios tectonic graben, which is considered to be an extension of the Anatolia strike-slip fault (Georgalas and Papakis, 1966, Marinos, Frangopoulos, & Stournaras, 1973). The activity of trending faults contributes to the uprise of thermal water, which specifically for the Thermopyles springs mixes with seawater or freshwater resulting in lower temperatures, compared to other geothermal springs, close to 40 °C. (Duriez et al., 2008). The system is rich in ammonia and hydrogen sulfide produced by the reduction of sulfate originating from the oxidation of sulfide minerals or directly from seawater (Lambrakis, Katsanou, & Siavalas, 2014).

Very few studies have been conducted in the Thermopyles and these were focused on geochemical features and measurements of physicochemical properties (Duriez et al., 2008; Lambrakis & Kallergis, 2005; Lambrakis et al., 2014; Verros et al., 2007; Zarikas et al., 2014) or the investigation of travertine deposits in association with *Cyanobacteria* (Kanellopoulos, Lamprinou, Mitropoulos, & Voudouris, 2016). Investigation of geochemical features has shown the increased concentration of hydrogen sulfide mainly produced by pyrite oxidation and reduced species of sulfates (Duriez et al., 2008; Lambrakis & Kallergis, 2005). The microbial community of Thermopyles has only been studied once (Kormas, Tamaki, Hanada, & Kamagata, 2009) but this previous study was focused on a cross-sectional comparison of populations to those in springs in other parts of Greece and was based on clone libraries that provided only coarse resolution.

Here we analyze shotgun metagenomes from 17 Thermopyles samples collected during a seven-year period and in different seasons along with environmental data in order to evaluate i) potential seasonal changes in microbial diversity and function, ii) the novelty of the microbial species that are always prevalent in the community using 16S rRNA and metagenome assembled genomes (MAGs), and iii) key metabolic functions that fuel the microbial communities in terms of energy and carbon sources for further biotechnological applications. Our findings suggested that seasonal and temporal changes were not significant in shaping microbial community composition and function while microbial communities were characterized by several novel species that have a key role in habitat function.

## MATERIALS AND METHODS

### Sample collection and processing

Samples were collected from the Thermopyles geothermal springs from June 2010 until December 2016 (Table S1). Water samples of 5 l were collected in pre-sterilized dark polyethylene bottles from a seepage point with visual water flow and bubbling. Upon return to the lab and within 3 hours, each sample was pre-filtered through a sterile 180 μm mesh nylon filter (Millipore, Burlington, MA, USA) to exclude sampling of microorganisms attached to large particles, and then a volume of 1.8 – 4.5 l of filtrate from the pre-filtration was filtered under mild vacuum (<100 mm Hg) through a 0.2 μm isopore polycarbonate filter (Sartorius, Göttingen, Germany) to collect microbial biomass. Subsequently, filters were folded aseptically, placed in sterile cryovials and stored at −80°C until further processed. *In situ* measurements, including temperature, pH and conductivity (Table S1), were taken by a portable multisensor instrument (WTW/Xylem, Rye Brook, NY, USA). Subsamples of 10-15 ml were fixed with 2% formaldehyde final concentration and kept at 4°C in the dark for cell counts. These subsamples were filtered to black Nuclepore filters (pore size of 0.2 μm) and stained with DAPI (4’,6-diamidino-2-phenylindole)(0.1 ug/ml). DNA extraction was performed with the MoBio Power Soil kit (MoBio Inc.Carlsbad, CA, USA) following its standard protocol. For all samples, libraries were prepared using the Illumina Nextera XT DNA library prep kit according to manufacturer’s instructions, except that the protocol was terminated after isolation of cleaned double stranded libraries. An equimolar mixture of the libraries was sequenced on an Illumina HiSEQ 2500 instrument (High Throughput Sequencing Core, Georgia Institute of Technology) for 300 cycles (2 × 150 bp paired-end rapid run). Adapter trimming and demultiplexing of sequenced samples was carried out by the instrument.

### Metagenomic read sequence trimming and assembly

Illumina reads were trimmed using a Q = 20 Phred quality score cut-off using SolexaQA ++(Cox, Peterson, & Biggs, 2010) and only trimmed reads longer than 50bp were considered for further analysis. Metagenomic reads were assembled using IDBA with default settings for metagenomes (Peng, Leung, Yiu, & Chin, 2012). Protein-coding genes were predicted from contigs longer than 500bp using MetaGeneMark.hmm with default parameters (Zhu, Lomsadze, & Borodovsky, 2010). Sequencing and assembly statistics for the metagenomic datasets are provided in Table 1. Metagenomic datasets have been deposited in NCBI SRA under the bioproject PRJNA611516.

**Table 1.** Metagenomic dataset statistics associated with Thermopyles dataset.

| **WGS dataset statistics** | **AP10** | **JN10** | **JN11** | **DE11** | **MY12** | **DE12** | **AU13** | **DE13** | **MY14** |
| --- | --- | --- | --- | --- | --- | --- | --- | --- | --- |
| **Total size (Gb)** | 3.47 | 3.31 | 3.54 | 2.24 | 2.37 | 3.17 | 3.02 | 3.84 | 2.82 |
| **# predicted genes** | 44,818 | 92,624 | 40,118 | 49,796 | 39,349 | 45,342 | 37,093 | 54,317 | 45,364 |
| **# contigs** | 12,590 | 23,409 | 9,720 | 17,517 | 11,372 | 11,625 | 11,178 | 16,985 | 10,063 |
| **Average contig length** | 2,867 | 3,277 | 3,512 | 2,319 | 2,842 | 3,242 | 2,611 | 2,566 | 3,802 |
| **Longer contig (bp)** | 186,023 | 250,868 | 138,677 | 104,928 | 186,610 | 239,545 | 204,647 | 240,475 | 240,302 |
| **Coverage (Nonpareil)** | 0.85 | 0.77 | 0.92 | 0.84 | 0.81 | 0.86 | 0.88 | 0.85 | 0.93 |
| **N50** | 3,691 | 4,791 | 5,851 | 2,521 | 3,874 | 4,658 | 3,132 | 3,067 | 7,823 |

| **WGS dataset statistics** | **DE14** | **FE15** | **AP15** | **JN15** | **DE15** | **MY16** | **OC16** | **DE16** |
| --- | --- | --- | --- | --- | --- | --- | --- | --- |
| **Total size (Gb)** | 3.35 | 1.70 | 3.34 | 3.24 | 3.28 | 4.11 | 2.71 | 3.48 |
| **# predicted genes** | 112,357 | 16,152 | 31,367 | 102,440 | 61,816 | 80,003 | 35,720 | 29,561 |
| **# contigs** | 21,625 | 4,931 | 8,295 | 24,137 | 14,613 | 22,072 | 8,128 | 8,011 |
| **Average contig length** | 4,693 | 2,591 | 3,118 | 3,603 | 3,519 | 2,988 | 3,731 | 2,994 |
| **Longer contig (bp)** | 260,721 | 240,237 | 229,951 | 269,767 | 240,362 | 240,890 | 691,025 | 164,877 |
| **Coverage (Nonpareil)** | 0.86 | 0.95 | 0.85 | 0.88 | 0.92 | 0.87 | 0.94 | 0.93 |
| **N50** | 10,965 | 2,833 | 5,336 | 5,558 | 5,519 | 4,062 | 7,697 | 4,228 |

Metagenomic read encoding fragments of the 16S rRNA gene were identified with Parallel-META (v.2.1) (Su, Xu, & Ning, 2012), and were subsequently processed for OTU picking (>97% sequence identity threshold) and taxonomic identification with SILVA128 database (Pruesse et al.,2007) using MacQIIME 1.8.0 with default parameters (Caporaso et al.,2010).

### Population genome binning

Contigs longer than 500 bp were binned into MAGs using MaxBin v2.1.1 with default settings (Wu, Tang, Tringe, Simmons, & Singer, 2014). In each binning run, only contigs from the assembly of an individual sample were used (no co-assembly was performed). CheckM and the MiGA webserver (www.microbial-genomes.org) were used to estimate completeness and contamination of each MAG based on the recovery of single-copy universal bacterial proteins (Parks, Imelfort, Skennerton, Hugenholtz, & Tyson, 2015; Rodriguez-R et al., 2018). Recruitment plots and coverage for MAG contigs and genes were calculated using the ‘enveomics.R’ package v1.4.1 from the Enveomics Collection (Rodriguez-R & Konstantinidis, 2016). Final MAGs are available through: http://enve-omics.ce.gatech.edu/data/ and in NCBI SRA under the bioproject PRJNA611516. Relative abundance of MAGs was calculated as 100*MAG _coverage_*(MAG_size_(bp)/Metagenome _size_ (bp)). Taxonomy of the MAGs was estimated by MiGA with the NCBI Genome (Prokaryotes) and TypeMAT databases containing all quality controlled genomes from prokaryotic type material contained in NCBI until June 2020 (MiGA online; June 2020). Genomes were also classified using GTDB-tK in order to also include previously published MAGs for comparisons (Chaumeil, Mussig, Hugenholtz, & Parks, 2020). [MAGs naming scheme: the first part reflects the closest relative of the MAG and the second part the lowest taxonomic rank the two share according to the MiGA TypeMAT/NCBI database (p<0.1), i.e., C:class, O:order, F:family. G:Genus, S:Species. For instance, we use Sulfurimonas_O for a MAG that had a Sulfurimonas sp. as the closest relative and was classified - at the lowest level with statistical confidence- to the order C*ampylobacteriales*].

### Functional annotation of predicted genes and determination of differentially abundant features

The predicted protein sequences encoded in the MAGs and the assembled contigs were searched against uniprot-TrEMBL(2018) to assign functional annotation based on best matches, and ≥40% similarity, ≥60 bitscore and ≥ 80% alignment length cutoffs for a match. Predicted genes were further grouped in functional categories based on their best match against the SEED database using the subsystems categories (Overbeek et al., 2005) and the KEGG-orthology using Ghost-KOALA (Kanehisa, Sato, Kawashima, Furumichi, & Tanabe, 2016).

Metagenomic reads were mapped on the predicted genes from the assembly using BLAT (Kent, 2002) with at least 95% identity and 50% of query length aligned. The abundance of each gene on each dataset was estimated by the number of reads that mapped on the genes with the above cutoffs (gene coverage) after normalizing for gene length. The relative abundance of annotation terms (subsystems) in each dataset was estimated based on the sum of the abundances of the corresponding genes, normalized for dataset size (*i.e.*, total number of reads). Differentially abundant categories between samples were identified with the DESeq2 package version 3.0.2 (Anders & Huber, 2010) using the binomial test and false discovery rate <0.05.

### Correlation analysis

Spearman correlations between MAGs abundances and environmental factors were calculated using the PAST software (Hammer, Harper, & Ryan, 2001). Canonical Correspondence Analysis (CCA) between MAGs abundance and environmental parameters was performed using the Vegan package in R (Oksanen et al., 2013).

### Comparisons to other metagenomes

Available protein datasets from Kalamas River in Greece (kal2feb; Meziti, Tsementzi, Ar. Kormas, Karayanni, & Konstantinidis, 2016) and Yellowstone National Park (CIS_19; Inskeep et al., 2013) metagenomes were analyzed and annotated similarly to the Thermopyles datasets. These datasets were used in order to compare microbial community function between Thermopyles and another freshwater environment in Greece (Kalamas) and a geothermal spring in the United States (Yellowstone). Comparisons between metagenomic datasets were performed based on the gene counts of the different functional annotation terms (subsystems). To make these counts comparable between datasets, protein sequences were first clustered using the CD-HIT algorithm (Fu, Niu, Zhu, Wu, & Li, 2012) with the following parameters: S=97 (similarity threshold) and aL=0.5 (minimum length coverage). Representative proteins from each cluster were annotated based on their best match against the SEED database. Comparisons between gene counts of different subsystems were performed using DESeq2 as described above.

## RESULTS

### Microbial community structure of Thermopyles

In the metagenomic datasets, a total of 887,602 individual reads encoding fragments of the 16S rRNA gene were detected (Table S1). [Sample naming scheme: the two letters and the two numbers reflect the month and the year, in which the sample was collected. e. g. AU13 represents a sample collected in August 2013]. Bacterial sequences predominated since only 0.05% of the total reads were assigned to *Archaea*. The highest diversity as indicated by OTU richness measurements using Chao and Shannon indices was observed in the DE15 sample while the lowest value was observed for DE16 (Table S1). Good’s coverage values exceeded 0.98 samples indicating that approximately 98% of the total 16S rRNA gene-based OTUs were recovered by sequencing. Consistent with this finding based on 16S rRNA gene fragments, the majority (60.9-93.5%) of the metagenomic reads assembled in contigs longer than 500bp (Table S1). The coverage of the microbial community achieved by the corresponding metagenomic dataset was also estimated based on the redundancy of the reads using the Nonpareil algorithm (Rodriguez-R & Konstantinidis, 2014), which confirmed the relatively low complexity of all of the samples as evidenced by coverage values ranging from 0.77 to 0.94 (Table 1). In total 5,956 OTUs were observed in all time points. Only 66% of these OTUs had a >97% identity match against the SILVA database, while the rest represented ‘novel’ OTUs (Table S1). Those novel OTUs comprised a substantial portion of the total community, accounting for 16%-40% of the total reads in each sample (Table S1).

Taxonomic composition was estimated based on the short reads encoding 16S rRNA gene fragments recovered in the metagenomes. *Proteobacteria* were the most abundant phylum across all samples (Fig. S1a). Class level taxonomic distributions revealed the dominance of *Epsilonproteobacteria* (>60% of total sequences) in most samples, usually followed by *Gamma-* and/or *Alphaproteobacteria* (Fig. S1b). [Note that the classification of *Epsilonproteobacteria* has been the target of several recent studies and the proposal of a separate phylum, *Epsilonbacteraeota*, has been suggested (Waite et al., 2017). Since then different databases (e.g., SILVA) have followed this suggestion while others have not (e.g., GenBank). In this study, different databases are used for classification, but we chose to use the GenBank classification of *Epsiloproteobacteria* as a Class within *Proteobacteria* phylum for consistency with most previous studies].

From the 5,956 OTUs observed in all time points, 165 represented shared OTUs, comprising 18.14-91.64% (average 74.09%) of the total 16S rRNA gene carrying reads per sample (Table S1). Given the similar sequencing depth across the samples (Table 1, S1), the comparison of the number of detected shared OTUs can provide a reliable picture of the size and composition of the “core” bacterial community in the springs. This ‘core’ bacterial community mainly consisted of representatives of the *Piscirickettsiaceae, Sulfurovaceae,* and *Campylobacteraceae* families, and the *Halothiobacillus, Sulfurovum, Arcobacter, Sulfuricurvum* and *Sulfurimonas* genera (Fig. S1c). Changes in the abundance of the core microbial communities did not exhibit any apparent seasonal pattern, which was consistent with the results of the Morisita analysis showing increased similarities between samples over long periods of time i.e. JN11 vs DE16 or JN10 vs JN15, interrupted by highly differentiated samples (DE11, DE14) (Fig. 1a). Apart from similarities in terms of taxonomic diversity, the size of the microbial community remained stable during the seven years as evidenced by DAPI counts ranging from 85,434 to 98,888 cells/ml (Table S1).

**Figure 1.**
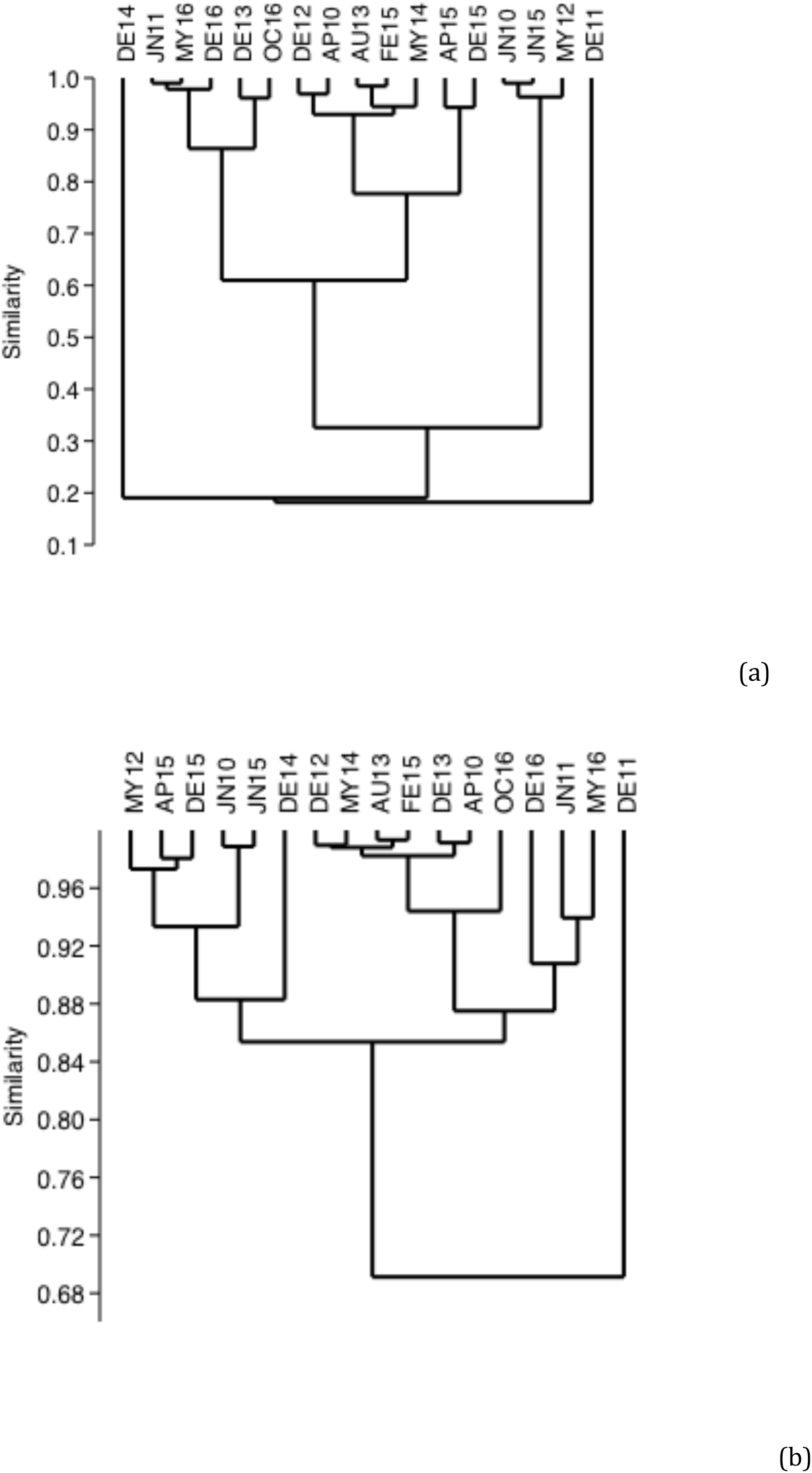
Clustering of Thermopyles samples based on a) OTU and b) seed subsystems relative abundance profiles

Consistent with the results reported above based on individual OTUs, our non-metric multidimensional scaling (NMDS) (stress: 12.57%) analysis performed at the whole community level based on the normalized abundances for metagenome size of all identified OTUs revealed no measured environmental parameters (e.g., season, precipitation, pH) to be significant for the ordination of the samples (Fig. S2). Further, cluster analysis revealed four distinct clusters exhibiting intra-cluster similarities exceeding 75% (vs. 32-60% between clusters), and two samples (DE11 and DE14) with much lower similarities to the rest of the samples (Fig. 1a). DE11 and DE14 showed <20% similarity to any other sample and to each other.

### Microbial functional diversity and comparisons to other habitats

The number of predicted genes from the metagenomic assemblies ranged from 29,561 (DE16) to 112,357 (DE14) (Table 1) reflecting the underlying microbial community diversity of the samples. Average contig length ranged from 2,318 bp (DE11) to 4,692 bp (DE14), while the largest contigs observed ranged from 104,928 bp (DE11) to 691,025 bp (OC16), (Table 1).

Functional distributions were evaluated based on the classification of genes in the subsystems hierarchical annotation scheme of the SEED database, resulting in 13.94% (DE11) to 54.16% (AG13) of total reads mapping on assembled genes that have SEED subsystems annotations (Table S1). Cluster analysis performed on the abundance distributions of different functions revealed higher Morisita similarities than the taxonomic (OTUs) similarities mentioned above, suggesting a greater number of shared functions among all samples (Fig. 1b). Consistent with the taxonomic results, DE11 exhibited the lowest functional similarities with the rest of the samples (<70%), while DE14 was more similar with the rest of the samples compared to the taxonomic diversity comparisons described above (Fig. 1b). Overall, DE11 exhibited higher abundances of CO_2_ fixation, metabolism of aromatic compounds, ammonia assimilation, phages prophages and transposable elements, sulfate reduction and CRISPRs related genes, and relatively lower abundance of denitrification, dissimilatory nitrite reduction and nitrate/nitrite ammonification compared to the other metagenomes (Fig. 2a). Pairwise comparisons using DeSeq2 of the functional distributions from all Thermopyles samples revealed relatively similar gene content with none of the 1,114 identified subsystems exhibiting significantly different abundances (p_adj_>0.05). In general, differentially abundant functions did not exhibit any specific pattern regarding season or month, similar to the OTU patterns mentioned above.

**Figure 2.**
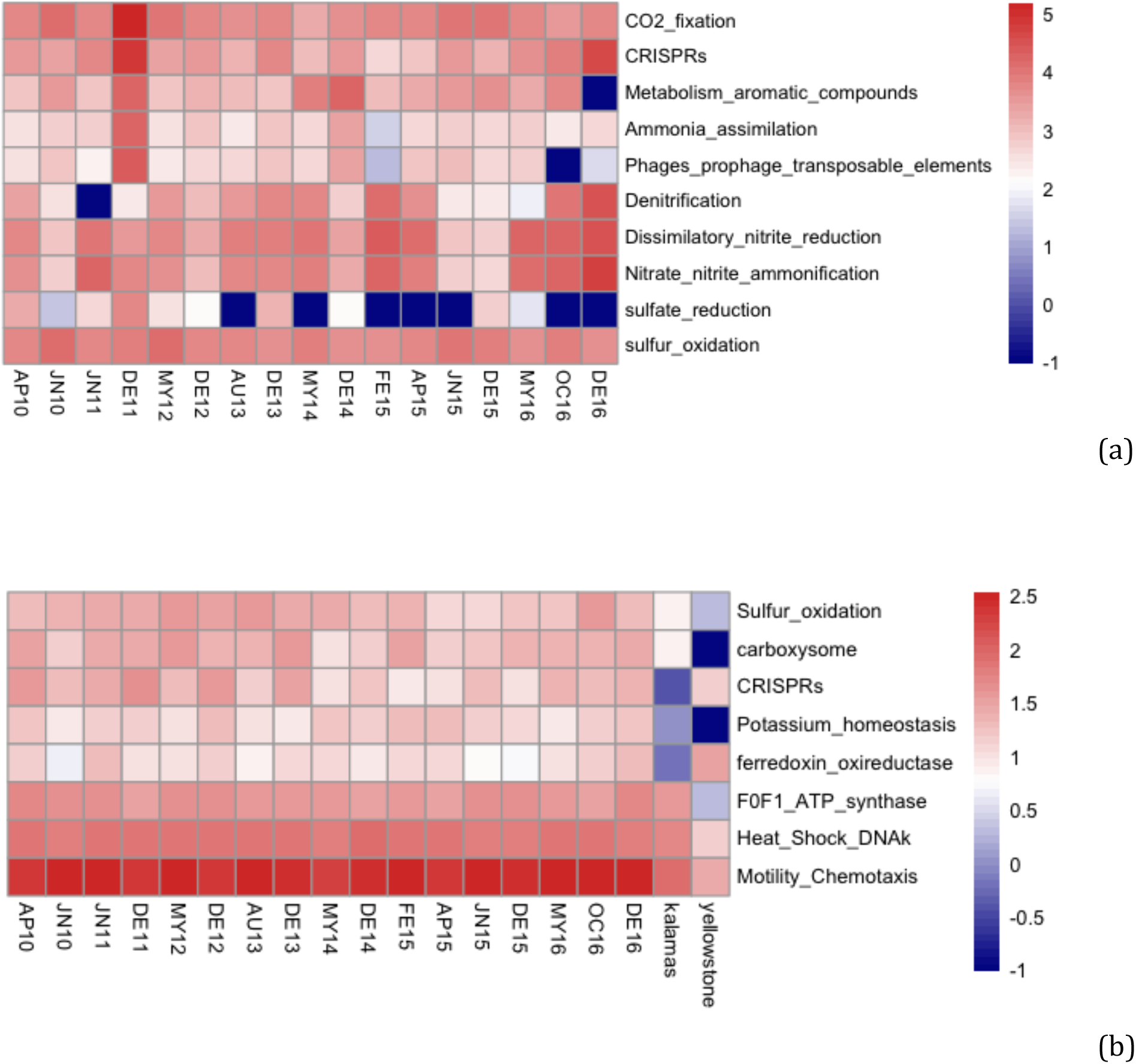
Functional profiles of Thermopyles microbial communities a) Subsystems (SEED database) with significant differences in abundance through time (Negative binomial test, DeSEQ, p<0.05; scale corresponds to number of normalized reads) b) Statistically significant differences in gene content (derived from the number of different genes that could be assigned to a subsystem) between Thermopyles, Yellowstone and Kalamas samples (Negative binomial test, DeSEQ,p_adj_<0.05; scale corresponds to normalized number of different genes).

When comparisons were performed with other habitats, several significant differences (p_adj_<0.05) were observed (Table S2, Fig. 2b) in terms of functional gene diversity. Overall, Thermopyles samples exhibited increased allelic diversity (i.e., a higher number of distinct variants of the same protein function at the 97% amino-acid similarity threshold) compared to both Kalamas and Yellowstone for all genes involved in sulfur oxidation pathways including all genes from the *sox* pathway as well as genes for flagellar motility, potassium homeostasis and the carboxysome. Flagellar motility genes are generally increased in springs relative to riverine samples since microorganisms use motility in order to find the niche with the most favorable concentrations of oxygen and nutrients. The relative lower frequency of flagellar motility genes found in the Yellowstone samples could be attributed to the decreased water flow (e.g., samples collected mostly from bottom or suspended sediment) in this ecosystem vs. the Thermopyles one. Similarly the highly abundant carboxysome genes in Thermopyles were most probably associated with *Cyanobacteria* that are abundant in the travertine deposits of Thermopyles (Kanellopoulos et al., 2016) but are absent from the specific site studied in Yellowstone. Gene diversity was increased in both Thermopyles and Yellowstone compared to Kalamas for CRISPR genes as well as ferredoxin oxidoreductase homologs, which are important in the reverse TCA cycle often observed in sulfidic and anaerobic environments dominated by *Epsilonproteobacteria* and *Archaea* (Fig. 2b). Diversity in CRISPR related genes was much higher in Thermopyles where 160 different genes were detected in total including genes related to cas1-9, Cmr3/4/6, and Csd1/2/5/7 gene families while in Kalamas only the Cas2 related proteins were detected.

Sulfur and nitrogen metabolism are of special interest for the microbial ecology of Thermopyles due to the chemical composition of the groundwater feed, and thus the corresponding pathways were examined in more detail using KEGG annotations. For sulfur metabolism, three pathways were assessed, i.e., M00595 for thiosulfate oxidation by the *sox* pathway (*soxABCDXYZ*), M00596 for dissimilatory sulfate reduction (*dsr*) and M00176 for assimilatory sulfate reduction (*asr*). Most samples contained all genes of the *sox* pathway and partial detection was observed in the rest, similar to the *asr* pathway (Fig. 2C). The complete *dsr* pathway was detected in eight samples while partial detection was observed in three samples and complete absence in the rest. For the samples in which partial detection was observed the calculation of genome equivalents showed that absence was probably due to low coverage rather than real absence (Table S6). Genome equivalent analysis also showed higher abundance (3 to 50 fold) of *soxCDY* genes relative to the rest of the genes of the pathway, implying that members of the community could follow an alternative pathway for sulfur oxidation with the absence of *soxABZX* as suggested previously for other systems (Lahme et al., 2019).

Regarding nitrogen metabolism, KEGG pathways predicted to be present in Thermopyles samples included nitrogen fixation (*nif*; KEGG ID M00175), assimilatory nitrate reduction (*anr*; KEGG ID M00531), as well as dissimilatory nitrate reduction (dnr; KEGG ID M00530) pathways. Nitrogen fixation genes, *nifHDK* and *vnfH* were present in all metagenomes apart from FE15. The *dnr* pathway was complete in all metagenomes and *anr* was complete in all metagenomes apart from MY14 and DE14 in which gene *nirA* for the transformation of nitrite to ammonia was absent (Fig 3c).

**Figure 3.**
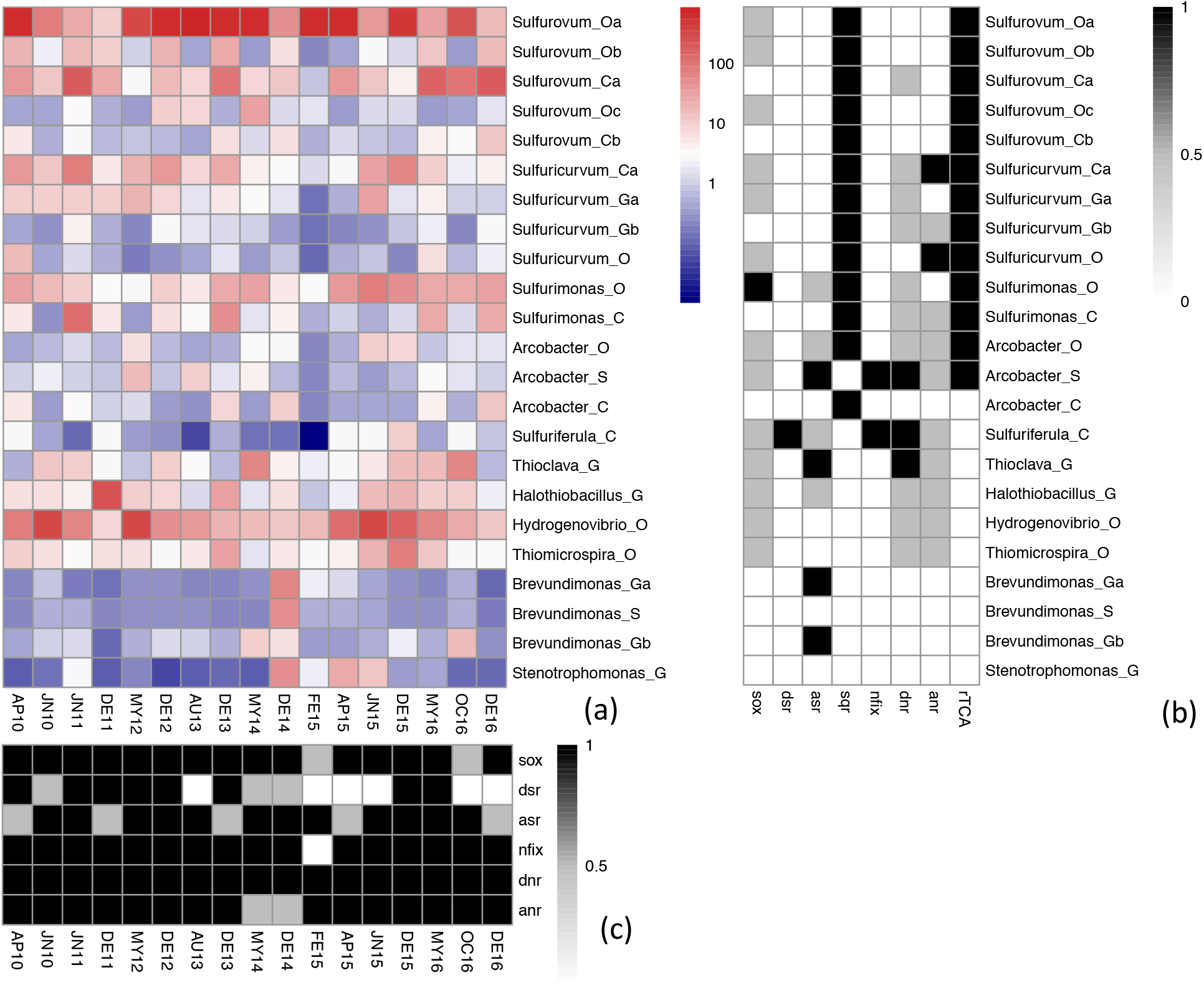
Abundance of MAGs and presence of specific pathways in metagenomes and MAGs a) Abundance of the identified bacterial populations (MAGs); scale: coverage. b) Presence absence of specific pathways in different MAGs c) Presence/absence of specific pathways in different metagenomes *dsr*: dissimilatory sulfate reduction, *asr*: assimilatory sulfate reduction, *sqr*: sulfide oxidoreductase, *nifix*: nitrogen fixation, *dnr*: dissimilatory nitrate reduction, *anr*: assimilatory nitrate reduction, denitr: denitrification, rTCA: reductive TCA cycle.

Amongst the total SEED subsystems detected in at least one of the samples, 250 out of 1,234 were common in all the habitats analyzed, and were present in more than half (eight) of the Thermopyles samples. Genes coding for these proteins were considered orthologous and were further analyzed for their %G+C content. The %G+C content in Thermopyles was significantly higher (61.6%) than the respective %G+C contents in Kalamas (47.2%) and Yellowstone (55.1%) (t-test, p<0.0001).

### Novelty of the Metagenome Assembled Genomes (MAGs)

In total, 78 good quality MAGs, i.e., completeness >70% and contamination (<10%), were recovered from the 17 samples. After ANI pairwise comparisons, the MAGs were grouped using a cutoff of 95% (threshold for species) in 43 genomospecies (GSP) for further analysis (Table 2). The best quality (i.e., highest completeness, lowest contamination) MAG of each genomospecies was used as a representative.

**Table 2.** Taxonomic identification of MAGs and assembly statistics. Best taxonomic classification (p<0.1) and closest relative according to MiGA NCBIprok and GTDB-tk databases.

| Sample | Bin | MAG name | Taxonomy NCBI (p<0.10) | closest_relative NCBI (AAI) | Taxonomy (GTDB-tk) | closest_rel GTDB-tk | Completeness | Contamination | # genes | length (bp) | GC% | N50 |
| --- | --- | --- | --- | --- | --- | --- | --- | --- | --- | --- | --- | --- |
| DE13 | 14 | Arcobacter_C | *Epsilonproteobacteria* (Class) | Arcobacter sp. L NC 017192 (47.14% AAI) | Campylobacteriales | nd | 82.9 | 9.9 | 1637 | 1369554 | 34.74 | 3,397 |
| JN15 | 18 | Arcobacter_O | Campylobacteriales (order) | Arcobacter sp. L NC 017192 (55.41% AAI) | Campylobacteriales | nd | 94.6 | 9 | 4141 | 3329994 | 33.02 | 5,788 |
| AU13 | 5 | Arcobacter_S | *Arcobacter* sp. L (Species) | Arcobacter sp. L NC 017192 (86.86% AAI) | Aliarcobacter sp002452 | GCA_002452915 | 84.7 | 9.9 | 3030 | 2361362 | 28.07 | 3,008 |
| DE14 | 2 | Brevundimonas_Ga | *Brevundimonas* (genus) | Brevundimonas diminuta NZ CP021995 (81.06% AAI) | Brevundimonas diminuta | GCA_002430835 | 75.7 | 1.8 | 3446 | 3497516 | 66.88 | 46,130 |
| OC16 | 6 | Brevundimonas_Gb | *Brevundimonas* (genus) | Brevundimonas sp. DS20 NZ CP012897 (83.75% AAI) | Brevundimonas | GCA_003248925.1 | 93.7 | 6.3 | 3844 | 3538981 | 68.25 | 34,665 |
| DE14 | 3 | Brevundimonas_S | *Brevundimonas* diminuta | Brevundimonas diminuta NZ CP021995(96.83) | Brevundimonas diminuta | GCF_000204035 | 73 | 1.8 | 3439 | 3418976 | 67.76 | 51,630 |
| JN15 | 14 | Halothiobacillus_G | *Halothiobacillus* (Genus) | Halothiobacillus neapolitanus c2 NC 013422T (89.46%) | Halothiobacillus | GCA_002282715 | 83.8 | 2.7 | 2684 | 2257048 | 56.06 | 4,350 |
| MY16 | 11 | Hydrogenovibrio_O | Thiotrichales (Order) | Hydrogenovibrio crunogenus XCL 2 NC 007520 (63.68% AAI) | Hydrogenovibrio | GCF_000711315.1 | 89.2 | 0.9 | 2345 | 2140471 | 45.1 | 8,061 |
| AP15 | 6 | Stenotrophomonas_G | *Stenotrophomonas* (genus) | Stenotrophomonas rhizophila NZ CP007597T (86.11% AAI) | Stenotrophomonas maltophilia | GCF_002799095 | 86.5 | 0.9 | 3624 | 4049585 | 67.72 | 56,024 |
| DE12 | 3 | Sulfuricurvum_C | *Epsilonproteobacteria* (Class) | Sulfuricurvum kujiense DSM 16994 NC 014762T (55.21% AAI) | Campylobacteriales | nd | 88.3 | 4.5 | 2470 | 2195547 | 45.61 | 9,117 |
| JN15 | 6 | Sulfuricurvum_Ga | *Sulfuricurvum* (Genus) | Sulfuricurvum kujiense DSM 16994 NC 014762T (79.16% AAI) | Sulfurimonadaceae | nd | 91 | 2.7 | 1787 | 1533918 | 45.58 | 9,298 |
| DE16 | 5 | Sulfuricurvum_Gb | *Sulfuricurvum* (Genus) | Sulfuricurvum kujiense DSM 16994 NC 014762T (79.05% AAI) | Sulfurimonadaceae | GCA_002633015.1 | 92.8 | 2.7 | 1693 | 1500498 | 44.39 | 10,849 |
| AP10 | 10 | Sulfuricurvum_O | Campylobacteriales (order) | Sulfuricurvum kujiense DSM 16994 NC 014762T (55.56%) | Sulfurimonadaceae | nd | 72.1 | 0.9 | 1817 | 1606347 | 57.92 | 16,172 |
| DE15 | 11 | Sulfuriferula_C | *Betaproteobacteria* (Class) | Sulfuriferula sp. AH1 NZ CP021138 (53.45%) | Thiobacillus denitrificans | GCA_002343685 | 95.5 | 8.1 | 4449 | 3975807 | 65.8 | 34,175 |
| MY16 | 8 | Sulfurimonas_C | *Epsilonproteobacteria* (Class) | Sulfurimonas autotrophica DSM 16294 NC 014506T (54.43% AAI) | Sulfurimonadaceae | GCA_002633015 | 84.7 | 1.8 | 1645 | 1460674 | 44.24 | 8,986 |
| MY14 | 6 | Sulfurimonas_O | Campylobacteriales (order) | Sulfurimonas autotrophica DSM 16294 NC 014506T (64.07% AAI) | Sulfurimonadaceae | nd | 91 | 7.2 | 2287 | 1993344 | 37.39 | 10,100 |
| DE12 | 5 | Sulfurovum_Ca | *Epsilonproteobacteria* (Class) | Sulfurovum sp. NBC37 1 NC 009663 (56.17% AAI) | Sulfurovaceae | GCA_001493195 | 82 | 6.3 | 1913 | 1437704 | 34.37 | 3,088 |
| DE16 | 8 | Sulfurovum_Cb | *Epsilonproteobacteria* (Class) | Sulfurovum lithotrophicum NZ CP011308T (52.68% AAI) | Campylobacteriales | nd | 93.7 | 3.6 | 1914 | 1688626 | 33.98 | 11,871 |
| AP10 | 1 | Sulfurovum_Oa | Campylobacteriales (order) | Sulfurovum lithotrophicum NZ CP011308T (63.3% AAI) | Sulfurovaceae | GCA_002742845.1 | 92.8 | 4.5 | 1839 | 1765907 | 41.56 | 72,213 |
| AP10 | 8 | Sulfurovum_Ob | Campylobacteriales (order) | Sulfurovum lithotrophicum NZ CP011308T (56.37% AAI) | Sulfurovaceae | GCA_002311915.1 | 89.2 | 3.6 | 1861 | 1745349 | 42.47 | 7,244 |
| MY14 | 5 | Sulfurovum_Oc | Campylobacteriales (order) | Sulfurovum lithotrophicum NZ CP011308T (63.07% AAI) | Sulfurovaceae | GCF_001595645 | 93.7 | 1.8 | 2082 | 1968869 | 46.8 | 25,751 |
| JN11 | 13 | Thioclava_G | *Thioclava* (Genus) | Thioclava nitratireducens NZ CP019437T (82.65% AAI) | Thioclava marina | GCF_002020135 | 92.8 | 2.7 | 4007 | 3856190 | 64.05 | 19,975 |
| DE12 | 2 | Thiomicrospira_O | Thiotrichales (Order) | Thiomicrospira aerophila AL3 NZ CP007030T (56.15% AAI) | Thiomicrospiraceae | GCF_000483485 | 84.7 | 0.9 | 2296 | 2067329 | 51.78 | 5,551 |
| AU13 | 4 |  | Rhodobacteraceae (Family) | Rhodobacter sp. LPB0142 NZ CP017781 (68.25% AAI) |  |  | 73.9 | 2.7 | 3568 | 3413188 | 67.28 | 9,731 |
| AP15 | 5 |  | Sphingomonadales (Order) | Altererythrobacter mangrovi NZ CP022889T (62.74% AAI) |  |  | 92.8 | 9.9 | 3601 | 3597969 | 65.04 | 45,746 |
| DE14 | 4 |  | Micrococcales (order) | Cellulomonas fimi ATCC 484 NC 015514T (65.53% AAI) |  |  | 90.1 | 2.7 | 3778 | 4066437 | 75.38 | 16,325 |
| DE14 | 9 |  | Microbacterium (Genus) | Microbacterium aurum NZ CP018762(93.92) |  |  | 89.2 | 0.9 | 2757 | 2881951 | 70.61 | 44,540 |
| DE15 | 6 |  | Rhodobacteriales (Order) | Hyphomonas neptunium ATCC 15444 NC 008358T (61.45% AAI) |  |  | 92.8 | 1.8 | 3514 | 3552243 | 61.4 | 68,082 |
| DE15 | 12 |  | Micrococcales (order) | Phycococcus dokdonensis NZ LT629711T (61.57% AAI) |  |  | 83.8 | 1.8 | 4173 | 3855105 | 73.5 | 8,018 |
| JN10 | 3 |  | Rhodobacteriales (Order) | Rhodobacter blasticus NZ CP020470 (62.39% AAI) |  |  | 91.9 | 0.9 | 3413 | 3182379 | 70.24 | 8,962 |
| JN10 | 6 |  | *Gammaproteobacteria* (Class) | Gallaeimonas mangrovi NZ CP031416T (45.28% AAI) |  |  | 95.5 | 0.9 | 2268 | 2169198 | 46.34 | 23,813 |
| JN10 | 7 |  | *Rhizorhabdus* (Genus) | Rhizorhabdus dicambivorans NZ CP023449(81.5) |  |  | 95.5 | 1.8 | 4109 | 4151982 | 65.25 | 68,680 |
| JN11 | 10 |  | *Stenotrophomonas* (genus) | Stenotrophomonas maltophilia NZ CP011305 (95.54) |  |  | 95.5 | 1.8 | 4489 | 4891136 | 66.09 | 51,010 |
| JN15 | 4 |  | Sphingobacteriales (Order) | Pseudopedobacter saltans DSM 12145 NC 015177T (63.25% AAI) |  |  | 95.5 | 0.9 | 3062 | 3278549 | 35.6 | 91,434 |
| JN15 | 10 |  | *Sphingopyxis terrae* (Species) | Sphingopyxis terrae subsp. terrae NBRC 15098 CP0133429(94.12) |  |  | 87.4 | 4.5 | 3293 | 3372237 | 65.39 | 83,053 |
| JN15 | 11 |  | Sphingomonadales (Order) | Altererythrobacter mangrovi NZ CP022889T (62.98% AAI) |  |  | 73 | 0.9 | 3061 | 3068126 | 65.71 | 51,503 |
| JN15 | 15 |  | *Sphingopyxis* (Genus) | Sphingopyxis macrogoltabida NZ CP009429T (81.56% AAI) |  |  | 79.3 | 3.6 | 3787 | 3833590 | 65.76 | 33,596 |
| JN15 | 21 |  | Micrococcales (order) | Microbacterium chocolatum NZ CP015810 (66.76% AAI) |  |  | 80.2 | 1.8 | 3430 | 2879452 | 66.83 | 4,351 |
| MY12 | 9 |  | Moraxella osloensis (Species) | Moraxella osloensis NZ AP017381 (95.4) |  |  | 89.2 | 6.3 | 2774 | 2538488 | 44.02 | 4,340 |
| MY14 | 4 |  | Flavobacteriaceae (Family) | Flavobacterium sediminis NZ CP029463T (74.85% AAI) |  |  | 73.9 | 0.9 | 2497 | 2179174 | 30.65 | 7,369 |
| MY14 | 9 |  | *Gammaproteobacteria* (Class) | Aeromonas schubertii NZ CP013067 (46.72% AAI) |  |  | 88.3 | 2.7 | 2632 | 2499300 | 52.55 | 8,814 |
| OC16 | 5 |  | *Pseudomonas balearica* (Species) | Pseudomonas balearica DSM 6083 NZ CP007511 (98.220) |  |  | 94.6 | 1.8 | 4283 | 4536722 | 64.76 | 196,460 |

Our assessment revealed that the majority of the MAGs (15/43) belonged to *Epsilonproteobacteria* followed by *Gammaproteobacteria* (Table 2), consistent with the 16S rRNA gene-based OTU results mentioned above. The majority of the *Epsilonproteobacteria* affiliated MAGs were only classified to the order or class level compared to all previously named species of isolates, suggesting that they represent novel families, not previously classified and/or with no genome representatives. Results from GTDB-tK classifications for the *Epsilonproteobacteria* MAGs were consistent, showing that our MAGs probably represent novel genera and/or families (Table 2). This finding partially agreed with the 16S results that showed ~15% of the *epsilonproteobacterial* classified OTUs belong to novel genera and 8% of them to novel families (Table S1).

In order to better study the taxonomy and function of abundant MAGs, 23 MAGs exhibiting high relative abundance (Table S3) and potential involvement in the sulfur cycle based on their predicted gene content were chosen for further investigation (Table 2, Fig.3a). Five of these MAGs (Sulfurovum_X) belonged to closely related species exhibiting ANI similarities below 95% but above 72% among them (Table S4). These MAGs had *Sulfurovum lithotrophicum* NZ CP011308 (Jeon et al., 2015) as a closest relative exhibiting AAI similarities between 52.68%-63.05% based on MiGA (Table 2); thus, they share only class and/or order with their closest relative and represent a novel genus, if not family. The *Sulfurovum* MAGs were dominant in most samples with coverage and relative abundances exceeding 100X and 20% of total community, respectively (Fig. 2A, Table S3).

The rest of the MAGs were taxonomically classified (p<0.1) to *Epsilonproteobacteria* or *Campylobacterales* having *Sulfuricurvum*, *Arcobacter* and *Sulfurimonas* genera as best matches with closest relatives *Sulfuricurvum kujiense* DSM 16994 NC 014762T, *Arcobacter* sp. L NC 017192 and *Sulfurimonas autotrophica* DSM 16294 NC 014506T, respectively; Table 2). ANI values between MAGs with the same closest relative varied from 71.14% to 90.32%, implying that they most likely belong to the same family or genus and revealed substantial intra-genus diversity at Thermopyles similar to the *Sulfurovum* MAGs mentioned above (Table S4). The remaining MAGs were taxonomically classified to *Brevundimonas*, *Halothiobacillus*, *Thioclava*, *Stenotrophomonas*, *Thiotrhichales* and *Betaproteobacteria.*

### Key Metabolic Functions of the MAGs

Key energy generating pathways were searched in the 23 abundant MAGs (Table 2) suspected to be involved in the sulfur cycle, including pathways for sulfur and nitrogen cycling as well as for carbon fixation. In addition to the KEGG pathways for sulfur and nitrogen cycling mentioned above, pathways for reductive TCA cycle (M00173) and sulfide quinone reductase (sqr; E.C.1.8.5.4) for the oxidation of sulfide to elemental sulfur were analyzed.

Using a similar approach and thresholds as mentioned above for assembled contigs and genes, *sqr* was detected in all *Epsilonproteobacteria* classified MAGs indicating the formation of elemental sulfur. Genes for the oxidation of thiosulfate by the sox pathway was partially observed in some *Sulfurovum, Sulfuricurvum*, *Arcobacter*, *Halothiobacillus* and *Thioclava* MAGs, while the complete complex was observed only in *Sulfurimonas*_O (Fig. 3b) agreeing with previous studies (Han & Perner, 2015). The absence of some genes from the *sox* complex could be attributed to low completeness in some cases such as with the *Sulfurovum* MAGs. However, the absence of specific genes (*soxCD*) in the *Sulfuricurvum* MAGs could indicate alternative pathways previously detected in *Sulfuricurvum* sp., in which thiosulfate oxidation ends up with the accumulation of sulfur globules or polysulfide (Friedrich, Bardischewsky, Rother, Quentmeier, & Fischer, 2005; Frigaard & Dahl, 2008).

Regarding *dsr*, only *Sulfuriferula*_C possessed the complete pathway while no related genes were observed in the other MAGs. For *asr*, the complete pathway was only detected in *Thioclava, Arcobacter*_S and *Brevundimonas*_Ga, while *Sulfurimonas*, *Arcobacter*, *Sulfuriferula*_C, *Halothiobacillus* and *Brevundimonas* MAGs possessed only some genes of the pathway mostly only including the genes for the first step, that is the transformation of sulfate to adenosine 5’ phosphosulfate (APS) (Fig.3b). Almost all of the MAGs possessed the enzyme sulfate adenyltransferase (*sat*), which is one of the enzymes responsible for the formation of APS in *dsr* and *asr* without possessing any other enzymes of the pathway. In these cases, both pathways were considered absent.

For nitrogen metabolism, the enzymes for nitrogen fixation were detected in *Arcobacter* and *Sulfuriferula* MAGs, along with the complete *dnr* pathway. The *dnr* pathway was also present in *Thioclava*_G and a partial pathway, missing the enzymes for production of nitrite, was present in *Sulfurovum*, *Sulfuricurvum* and *Sulfurimonas*. The remaining MAGs possessed only a few or no genes of the pathway. The *anr* pathway was complete only in *Sulfuricurvum* and genes for enzymes catalyzing the first step of anr (that is formation of nitrite) were detected in *Arcobacter*_S, *Thioclava*_G, *Sulfurimonas*_C and *Thiotrichales* MAGs (Fig.3b). These findings agreed with previous studies showing that nitrate can be used as an alternative electron acceptor for members of the *Sulfurovum, Sulfuricurvum, Sulfurimonas* and *Arcobacter* genera (Hamilton, Jones, Schaperdoth, & Macalady, 2015; Han & Perner. 2015).

Finally the TCA cycle was observed in all MAGs as expected but the key genes for the reductive TCA cycle, i.e., ATP citrate lyase; fumarate reductase and 2-oxoglutarate:ferredoxin oxidoreductase were observed only in *Sulfurovum*, *Sulfuricurvum Sulfurimonas* and *Arcobacter* MAGs (Fig. 3b) indicating that these species are capable of CO_2_ fixation.

### Correlation of MAG abundances with environmental parameters

In general, only a few and rather weak correlations were observed between MAG relative abundances and environmental parameters (Table S5). Most notably, pH value was positively correlated with abundance of *Sulfuricurvum*_G, temperature was negatively correlated with *Sulfurovum*_O and positively with *Sulfurimonas*_O abundances, while conductivity was positively correlated with *Halothiobacillus* abundance. Several correlations in abundances were observed between MAGs (Table S5) indicating possible synergistic or antagonistic interactions. Canonical Correspondence Analysis (CCA) confirmed the abovementioned results on differences in community composition characterizing samples DE11 and DE14 at the MAG level, while pH and conductivity emerged as important factors (P<0.05) for the ordination of the samples based on MAG abundances (Fig. S3)

## Discussion

As mentioned above, Thermopyles are characterized by high sulfide concentrations; hence it is expected that sulfides are an important energy source for autotrophic microorganisms (Porter, Engel, Kane, & Kinkle, 2009; Han et al., 2012; Rossmassler, Hanson, & Campbell, 2016). Moreover, *Epsilonproteobacteria* have been detected to be key players in sulfur metabolism in sulfide rich environments (Hamilton et al., 2015; Han et al., 2012; Rossmassler, Hanson, & Campbell, 2016) and were also prevalent in the Thermopyles samples collected in 2005 (Kormas et al., 2009). In agreement with these expectations and previous results, *Epsilonproteobacteria* were prevalent in our time series and their relative abundances were largely unaffected by environmental factors as revealed by 16S rRNA gene and MAGs data (Fig.3a, S2). Further, the presence of ‘core’ species revealed that Thermopyles microbial communities are dominated by specific microbial populations, which probably interact among each other (synergistically or competitively) and represent mostly novel genera, if not families. Results from a previous study in Yellowstone Park agreed with the presence of core communities despite the latter study including only three samples collected for 3 years on a yearly basis and lacking seasonal resolution (De León, Gerlach, Peyton, & Fields, 2013).

Although it was expected that sulfur cycling related pathways would be prevalent in Thermopyles, our study represents the first actual documentation of prevalent microbially-mediated processes related to sulfur and thiosulfate oxidation as well as assimilatory and dissimilatory sulfate reduction in a terrestrial sulfur geothermal spring. Our analysis also revealed substantially higher functional than taxonomic similarity (Fig. 1), revealing core functions dominating Thermopyles regardless of the prevailing environmental conditions (Fig. 2a). Prevalent functions in Thermopyles compared to another (non-sulfur rich, non high temperature) freshwater habitat included CRISPRs, sulfur oxidation and reductive TCA cycle related genes (Fig. 2b). These results also agreed with previous results from similar habitats (López-López et al., 2013; Sikorski et al., 2010). Increased diversity of CRISPR related genes, representing different Cas, Cmr and Csc proteins, has also been detected in Yumthang geothermal spring system that exhibits similar physicochemical properties with Thermopyles (Najar, Sherpa, Das, & Thakur, 2020), CRISPR systems are considered stress regulators since they represent defensive tools by attacking viruses and plasmids (Louwen, Staals, Endtz, Baarlen, & Oost, 2014) and are more abundant in thermophilic than mesophilic bacteria, as has been shown in some environmental (Makarova, Grishin, Shabalina, Wolf, & Koonin, 2006; Westra, van Houte, Gandon, & Whitaker, 2019) and predictive modeling studies (Weissman, Laljani, Fagan, & Johnson, 2019).

The prevalence of proteins related to flagellar motility corroborates with results from other geothermal habitats (Badhai, Ghosh, & Das, 2015) but the lower abundances in Yellowstone sulfur rich sediment sample pointed out the importance of water flow characterizing different geothermal springs (Inskeep et al.,2013). Similarly the prevalence of carboxysome genes indicated the importance of *Cyanobacteria* in CO_2_ fixation at Thermopyles additionally to chemoautotrophy. This feature was absent from Yellowstone CIS_19 site, since this was an archaea dominated site, but other sites in Yellowstone, with temperatures and pH similar to Thermopyles, were mostly phototrophic and dominated from *Cyanobacteria.* Nonetheless, the latter sites lacked chemoautotrophy related pathways, showing once more the importance of different physical (i.e. flow) and geochemical (i.e. dissolved sulfides, temperature) characteristics, which, in combination, drive microbial community structure and metabolic properties.

Also at the genome level, concurrent with previous data for sulfide rich habitats (Hamilton et al., 2015), all the *Epsilonproteobacteria* MAGs analyzed possessed pathways for sulfur oxidation and CO_2_ fixation via the reductive TCA cycle. The presence of complete or incomplete sulfur oxidation pathways in the genome of the close relatives of the MAGs reported here, coupled to the frequent co-presence of sulfide quinone reductase (*sqr*) genes, further corroborated the metabolic versatility of *Epsilonproteobacteria* (Friedrich et al., 2005; Frigaard and Dahl, 2008; Hamilton et al., 2015,Han & Perner, 2015, Rossmassler et al. 2016, Wright, Williamson, Grasby, Spear, & Templeton, 2013).

The presence of different pathways oxidizing different sulfur compounds and producing either sulfate or elemental sulfur, along with correlation analysis results (Table S5, Fig. S3), implied that synergistic or antagonistic relationships between *Epsilonproteobacterial* populations could influence taxonomic and functional diversity at Thermopyles. Another explanation for the high intra-genus species and pathway diversity could be the ability to utilize alternative sulfur oxidation pathways when conditions are unfavorable or different affinities of different alleles for the same substrate exist. The importance of other environmental factors that were not measured here (i.e., metal concentrations) should be noted as a probable factor that could be included in future studies in order to better explain the prevalence of specific species at different time points.

Altogether, it appears that chemoautotrophic microbial communities that mainly oxidize different reduced sulfur compounds using oxygen or nitrate as electron acceptors inhabit Thermopyles. Although all sulfur oxidizing MAGs detected in this study probably consist of new genera that are members of a new family, they all belong to the *Epsilonproteobacteria* phylum, closely or remotely related to the genera *Sulfurovum, Sulfuricurvum, Sulfurimonas* and *Arcobacter* (Table 2). Similar communities have been observed in sulfur rich environments in the past (Hamilton et al.. 2015; Hotaling et al. 2019; Huegler et al., 2010; Rossmassler et al., 2016, Wright et al., 2013), but only once in a terrestrial geothermal spring (Reigstad et al., 2011) and cross-sectional (as opposed to a time series of seven years here) data. Notably, comparisons with genomes from some of these studies (when available) showed that the *Epsilonproteobacteria* detected in these previous studies, were remotely affiliated with the ones detected here. This indicated once more that Thermopyles prevalent bacteria are members of ‘novel’ genera and/or families that possess similar sulfur oxidizing properties with known species, further supporting a universal functional profile of *Epsilonproteobacteria* in sulfur rich environments.

The community composition of the two outlier samples with respect to stable core community (DE11 and DE14; Figs. 1a, 3a, S1c) was the only case that could be attributed to changes in environmental parameters such as conductivity and pH, respectively. From our data, it appears that changes in conductivity were positively correlated with precipitation during the days prior to sampling, though it has been previously noted that conductivity in Thermopyles is mostly influenced by sea tides that cause inflow of seawater in the spring (Zarikas et al., 2014). *Hallothiobacillus* MAG prevalence in DE11, along with its positive correlation with conductivity, was further supported by the known halotolerant character of *Halothibacillus* sp. not requiring salt in order to grow but growing optimally when salt concentrations increase (Sievert, Heidorn, & Kuever, 2000). Hence, we cannot conclude about the exact underlying cause of the unique diversity observed in samples DE11 and DE14, although it was most certainly related to inputs from non-geothermal sources.

The *Sulfuricurvum*_Ga MAG drop in abundance in MY14 and DE14 samples, along with a negative correlation with pH (Table S5), was further supported by previous studies showing that the type species of the genus, *Sulfuricurvum kujiense* (Kodama & Watanabe 2004), has a pH range between 6 and 8 while it grows optimally at pH 7 (Han et al., 2012). However, the prevalence of *Brevundimonas* related species in DE14 could not be linked to the pH drop since *Brevudimonas* species are alkalophiles, and this is further supported by their presence in alkaline thermal springs (Gupta, Gupta, Capalash, & Sharma, 2017; Tekere et al. 2011). The increased rainfall prior to sampling the DE14 sample, coupled with the 16S rRNA and MAG relative abundance and coverage data showing decreased *Epsilonproteobacteria* and increased *Alpha-, BetaProteobaceria* and *Actinobacteria* abundances might indicate influences from soil.

Finally, our results collectively showed that Thermopyles microbial communities are mainly fueled by chemolithotrophic processes performed by core sulfide oxidizing bacterial taxa that persist over time. Short intervals might be driven by changes in environmental parameters but changes in taxonomy never override the persistence of core functions. The outcome of this study, that is gene and genome sequences from ‘novel’ sulfur oxidizing bacteria, will be useful for the metagenomic investigation of other, similar environments. Future studies could assist on further understanding of evolutionary relationships and functionality of these and related microbial communities.

## Acknowledgements

This work was partly funded by the US National Science Foundation, awards #1831582 and #1759831 (to KTK), and by internal funding from the University of Thessaly (KAK). We thank the Partnership for an Advanced Computing Environment (PACE) at the Georgia Institute of Technology, which enabled the computational tasks associated with this study.

## Notes

### Competing Interest Statement

The authors have declared no competing interest.

