## Supplementary material for "Time series metagenomic sampling of the Thermopyles, Greece, geothermal springs reveals stable microbial communities dominated by novel sulfur-oxidizing chemoautotrophs": Suppl.Figures

Supplementary Figures

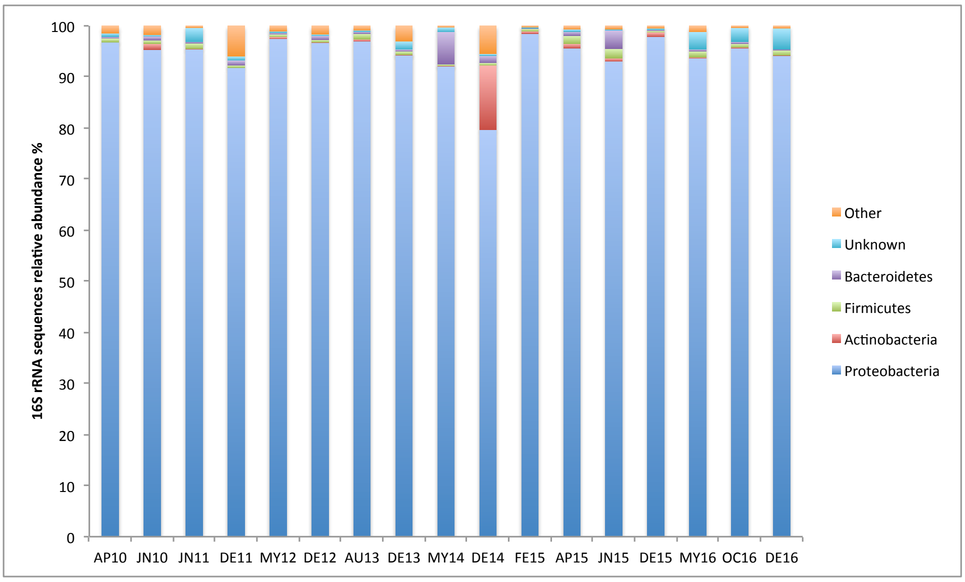

(a)

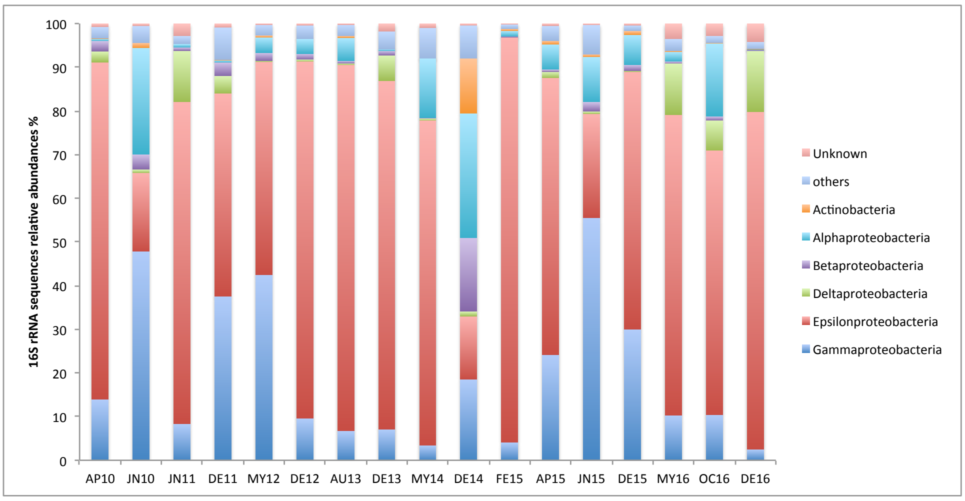

(b)

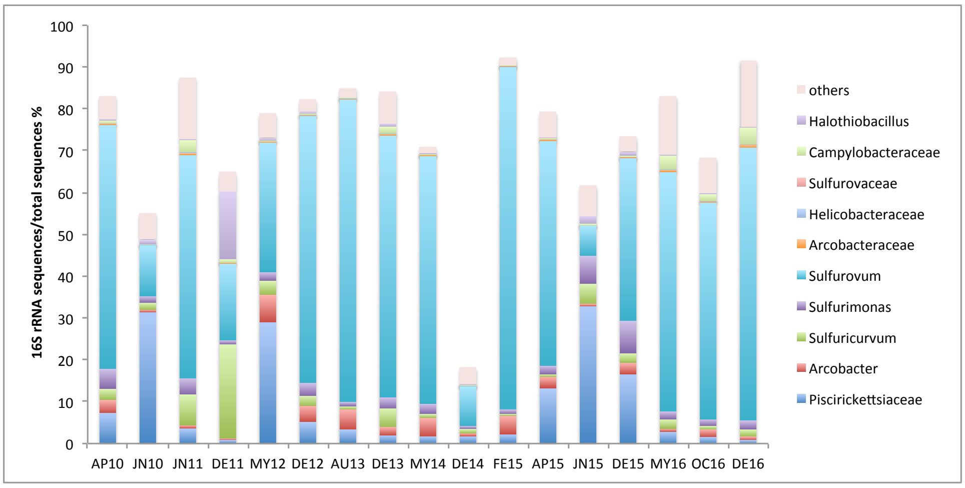

(c )

Figure S1. Taxonomic composition of Thermopyles geothermal springs microbial communities. Taxonomic distributions at the phylum (a) and class (b) level. Taxonomic distributions of 'core' OTUs.

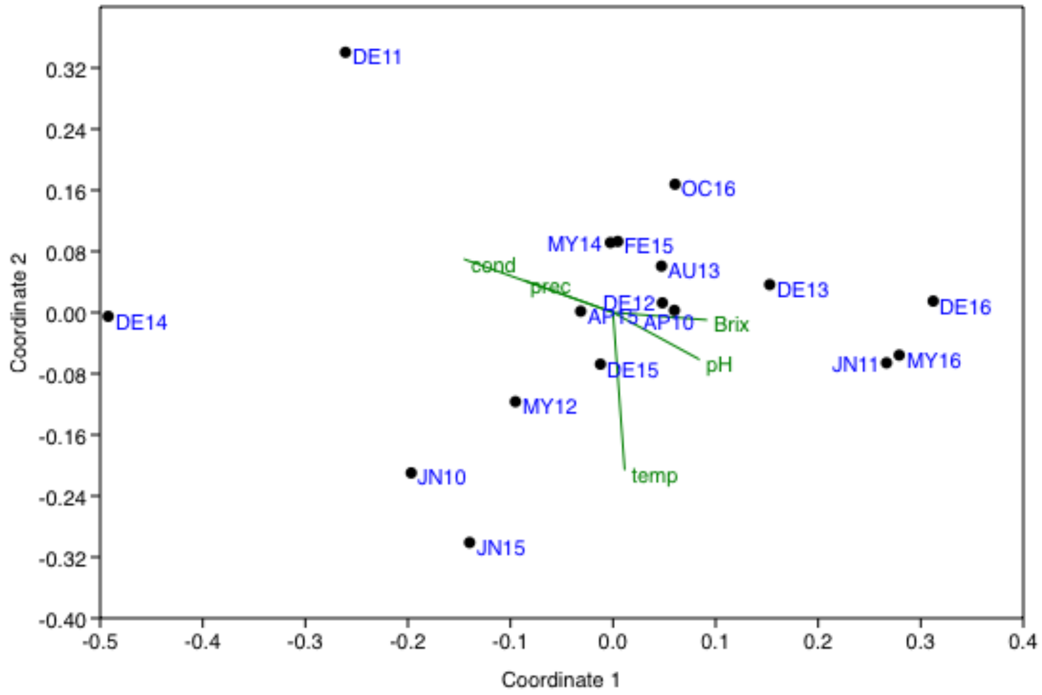


Figure S2. NMDS plot of taxonomic distributions based on abundances of OTUs (97% similarity) constructed from the metagenomes rRNA 16S gene-encoding reads.

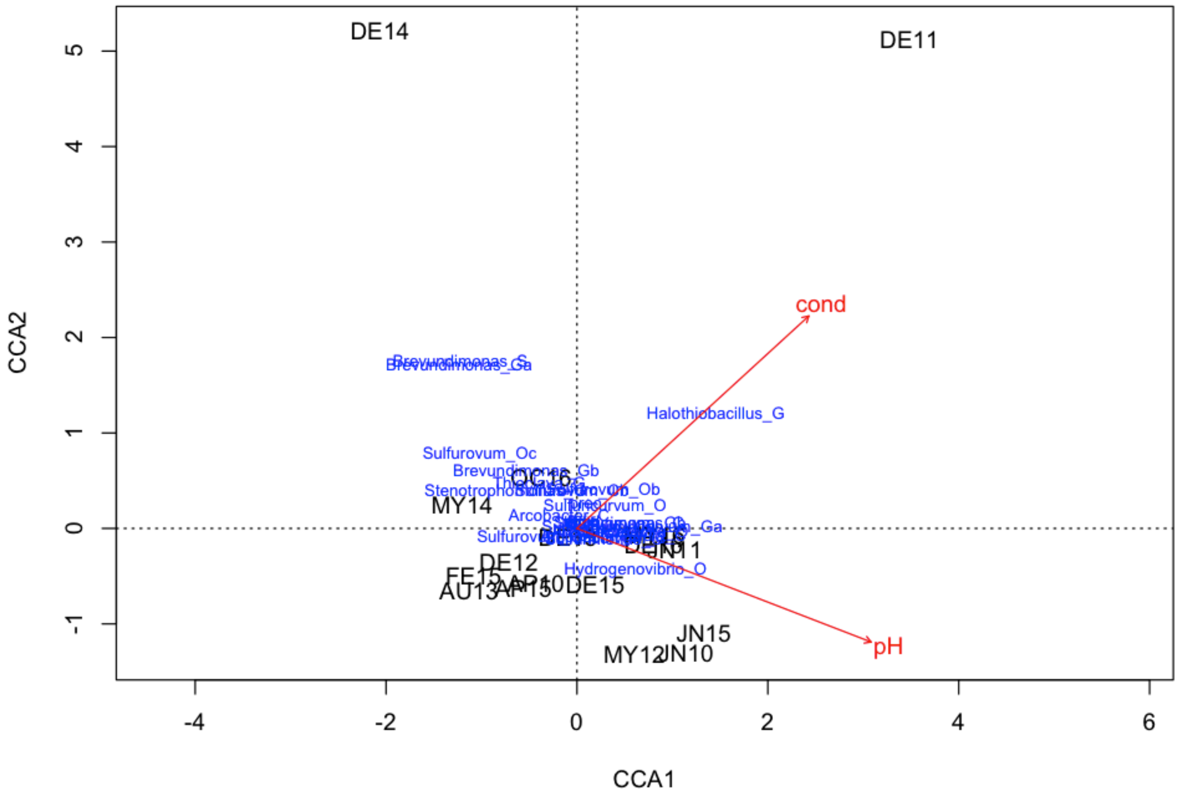


Figure S3. Canonical Correspondence Analysis (CCA) plot between MAGs relative abundance and environmental parameters.
